# Molecular signature of neutrophils in antiphospholipid syndrome based on epigenomic and transcriptome analysis

**DOI:** 10.1101/2025.10.03.680196

**Authors:** Moye Chen, Yujia Li

## Abstract

**Background:** Antiphospholipid syndrome (APS) is an autoimmune disorder characterized by thrombosis and pregnancy loss. Recent studies indicate that neutrophils, particularly neutrophil extracellular traps (NETs), contribute to the development and progression of APS. However, the precise underlying mechanisms remain unclear.

**Methods:** To address this, we analyzed epigenome and transcriptome data to identify key differentially expressed genes (DEGs) of neutrophils in APS using weighted gene co-expression network analysis (WGCNA). Two datasets (GSE102215 and GSE124565) were obtained from the GEO database. The Limma R package was used to identify DEGs, while ChAMP R was applied to analyze differentially methylated genes (DMGs) between APS patients and healthy controls. Enrichment analysis of Gene Ontology (GO) and Kyoto Encyclopedia of Genes and Genomes (KEGG) was performed using ClusterProfiler, and TISIDB was used to examine associations with immunomodulators, chemokines, and receptors. GeneCards and Connectivity Map databases were further used for disease-related gene analysis and drug prediction.

**Results:** GO enrichment analysis revealed that DEGs were primarily enriched in leukocyte cell–cell adhesion, regulation of leukocyte cell–cell adhesion, and cytokine-mediated signaling pathways. Correspondingly, KEGG enrichment analysis demonstrated that DEGs were mainly enriched in the ribosome, NF-kB signaling pathway, NOD-like receptor signaling pathway, and other related pathways. Through WGCNA analysis, we identified two key intersection genes, CCL5 and ITK, which were positively correlated with CD8+ T cells and eosinophils, while being negatively correlated with neutrophils and follicular helper T cells. Gene set enrichment analysis (GSEA) indicated that CCL5 was enriched in hematopoietic cell lineage, ribosome, and ribosome biogenesis in eukaryotes, and ITK was enriched in ribosome, ribosome biogenesis in eukaryotes, and T-cell receptor signaling. Similarly, gene set variation analysis (GSVA) showed that CCL5 and ITK were associated with IL-2–STAT5 signaling and the P53 pathway as well as MTORC1 signaling. Furthermore, gene regulatory network analysis suggested that CCL5 and ITK are modulated by common mechanisms involving multiple transcription factors. By examining APS-related genes in the GeneCards database, we observed a significant negative correlation between CCL5 and phosphatase and tensin homolog (PTEN) (R = −0.624), and a strong positive correlation between ITK and CD40LG (R = 0.933). Finally, drug perturbation expression profiles revealed that RS-56812, acebutolol, emetine, and proscillaridin exhibited the most significant negative correlation with disease-associated expression profiles.

**Conclusion:** These data nominate CCL5 and ITK as APS-associated genes in neutrophils and indicate associations with multiple signaling pathways. Moreover, drugs targeting these genes may represent potential therapeutic strategies for APS.

## Introduction

Antiphospholipid syndrome (APS) is an autoimmune disorder characterized by recurrent thrombosis and spontaneous abortion, both of which are strongly associated with the persistence of antiphospholipid (APL) antibodies (Handa, 2021). APS can occur independently as primary APS or in association with another autoimmune disease, such as systemic lupus erythematosus (SLE), in which case it is termed secondary APS (Tektonidou et al., 2004). Despite extensive research, the pathophysiology of APS remains incompletely understood. Previous work by Knight JS et al. demonstrated that neutrophils contribute to the pathogenesis of APS (Knight & Kanthi, 2022). Neutrophils from patients with APS are abnormally activated and display pro-inflammatory transcriptional potential (Qi et al., 2024). In particular, circulating APL has been shown to enhance reactive oxygen species (ROS) production by neutrophils (Salzer & Urban, 2024; Zaiema et al., 2024), thereby promoting the release of neutrophil extracellular traps (NETs). NETs are extracellular networks composed primarily of neutrophil-derived DNA and nucleosomes (Wang et al., 2024). Clinical studies have further revealed that patients with persistently positive APL often develop autoantibodies against NETs (Zuo et al., 2023). Interestingly, these antibodies against DNA and nucleosomes are also hallmark autoantibodies specific for SLE highlighting the immunological intersection between APS and SLE (Pisetsky, 2024).

Moreover, the presence of anti-NET antibodies impairs NET degradation leading to excessive accumulation of NETs in the circulation (Block et al., 2022). This excessive burden of NETs amplifies autoimmune inflammatory responses, enhances platelet activation, and drives thrombus formation (Perdomo & Leung, 2023), ultimately exacerbating the clinical manifestations of APS (Zhu et al., 2025). In addition to transcriptional abnormalities, neutrophils in APS also exhibit epigenetic alterations. Weeding E et al. reported that neutrophils from primary APS patients display 17 hypomethylated sites (including ETS1 and EMP2) and 25 hypermethylated CpG sites compared with healthy controls (Lopez-Pedrera et al., 2024). ETS1 has been shown to induce the differentiation of human villous trophoblast cells (Aplin & Jones, 2021), while EMP2 has been identified as a critical regulator of trophoblast function and vascular development in both mice and humans (Liu et al., 2024). Notably, EMP2-/-mice exhibit reduced litter size and placental hypoplasia, conditions linked to dysregulation of angiogenesis, coagulation, and oxidative stress pathways. These findings suggest that hypomethylation of ETS1 and EMP2 may contribute to APS-related fetal loss.

Despite these insights, research on neutrophils in APS at the gene level remains limited. To our knowledge, the present study is the first to employ a multi-omics weighted gene co-expression network analysis (WGCNA) using both transcriptome and epigenome microarray data to identify key neutrophil-associated genes in APS. Our aim is to explore how these genes contribute to disease development and progression, the signaling pathways in which they participate, and their potential as targets for gene-based therapeutic interventions.

## Materials

### 1. Data Acquisition and Preprocessing

GEO database (https://www.ncbi.nlm.nih.gov/geo/info/datasets.html) for GENE EXPRESSION OMNIBUS, is by the us national center for biotechnology information (NCBI database creation and maintenance of GENE EXPRESSION. The Series Matrix File of GSE102215 was downloaded from the NCBI GEO public database, and the annotated file was GPL16791. The expression profile data of 18 groups of samples were included, consisting of 9 healthy controls and APS patients. Similarly, the methylation Series Matrix File of GSE124565 was downloaded, with annotation file GPL13534. This dataset included 22 samples, comprising 12 healthy controls and 10 APS patients. Differential methylation sites were analyzed using the ChAMP package, with the screening condition set at adj.P.V < 0.05. In parallel, differential expression analysis was performed using the Limma package to identify DEGs between healthy controls and APS, with screening thresholds of P < 0.05 and logFC|> 0.585.

### 2. Function analysis of GO and KEGG

To investigate the biological functions and signaling pathways involved in APS, the R package “ClusterProfiler” was used to annotate the functions of DEGs. This allowed a comprehensive evaluation of their functional correlations. Both Gene Ontology (GO) and Kyoto Encyclopedia of Genes and Genomes (KEGG) analyses were conducted to assess functional categories. GO and KEGG enrichment pathways with p and q values < 0.05 were considered statistically significant.

### 3. WGCNA analysis

By constructing a weighted gene co-expression network, we can search for gene modules that are co-expressed, and explore the correlation between gene networks and phenotypes, as well as the key genes in the network. The coexpression networks of the genes in the dataset were constructed using the WGCNA-R package for further analysis, where the soft threshold was 3. The weighted adjacency matrix is transformed into topological overlap matrix (TOM) to estimate the network connectivity, and the hierarchical clustering method is used to construct the cluster tree structure of TOM matrix. Different branches of the cluster tree represent different gene modules, and different colors represent different modules. Based on the weighted correlation coefficient of genes, genes were classified according to their expression patterns, and genes with similar patterns were classified into one module, and tens of thousands of genes were divided into multiple modules by gene expression patterns.

### 4. GSEA analysis

GSEA was used to further analyze the differences of signaling pathways between high and low expression groups. Background gene sets were annotated gene sets of versions 7.0 downloaded from MsigDB database as annotation gene sets of subtype pathways. Differential expression analysis of pathways between subtypes was performed, and significantly enriched gene sets (adjusted p value less than 0.05) were sequenced according to consistency scores. GSEA analysis is often used to explore the close combination of disease classification and biological significance.

### 5. GSVA analysis (gene set difference analysis)

gene set variation analysis (GSVA) is a nonparametric, unsupervised method for evaluating transcriptome gene set enrichment. By comprehensively scoring the gene set of interest, GSVA converts the gene level change into pathway level change and then judges the biological function of the sample. In this study, gene sets were downloaded from Molecular signatures database (v7.0 version), and each gene set was comprehensively scored using GSVA algorithm to evaluate potential biological functional changes in different samples.

### 6. Regulatory network analysis of key genes

In this study, the R package “RcisTarget” was used to predict transcription factors. All calculations performed by RcisTarget are based on Motif. The normalized enrichment score (NES) for Motifs depends on the total number of Motifs in the database. In addition to the Motifs annotated by the source data, we infer further annotated files based on Motif similarity and gene sequences. The first step in estimating the overexpression of each Motif on the gene set is to calculate the area under the curve (AUC) of each pair of Motif-Motif sets. This is calculated by the recovery curve of the sequence sequence based on the gene set. The NES for each Motif was calculated from the AUC distribution of all Motifs in the gene set. We use rcistarge.hg19.motifdb.cisbpont.500bp for the Gene-Motif rankings database.Motif Enrichment Analysis Parameters.

### 7. miRNA network construction

miRNAs (MicroRNAs) are small non-coding RNAs that have been shown to regulate gene expression by promoting the degradation of mRNAs or inhibiting mRNAs translation. Therefore, we further analyzed whether there are some mirnas in key genes that regulate the transcription or degradation of some dangerous genes. We obtained key genes through TargetScanHuman database (https://www.targetscan.org/vert_80/) related micrornas, and through the cytoscape software visualization gene networks of micrornas.

### 8. Prediction of potential drugs

The Connectivity Map (CMap) is a gene expression profile database based on intervention gene expression developed by the Broad Institute. It is mainly used to reveal functional associations of small molecule compounds, genes, and disease states. It contains gene chip data before and after 1309 small molecule drugs are treated with five human cell lines. Treatment conditions are varied, including different drugs, different concentrations, different treatment duration, and so on. In this study, the differentially expressed genes of the disease were used to predict the targeted therapeutic drugs for the disease.

### 9. Statistical analysis

Statistical analysis: All statistical analyses were performed using R software (version 4.2.2). Unless otherwise specified, statistical tests were conducted as two-sided. An adj p-value < 0.05 was considered to indicate statistical significance. For gene expression analyses, multiple testing correction was applied where appropriate to control the false discovery rate. Correlation analyses were assessed using Spearman or Pearson methods depending on data distribution, and enrichment results were evaluated based on adjusted significance thresholds.

## Results

### 1. Integrated Transcriptomic and DNA Methylation Analyses of APS Cohorts

We first downloaded the dataset GSE102215 related to antiphospholipid syndrome from the GEO database, which included expression profile data from 18 patients, consisting of 9 healthy controls and 9 APS patients. The differential expression analysis was performed using the Limma package, with screening thresholds of P < 0.05 and |logFC| > 0.585. A total of 1,500 differentially expressed genes (DEGs) were identified, including 777 upregulated and 723 downregulated genes (**Fig. 1A–B**). To further investigate the biological relevance of these genes, pathway enrichment analysis was carried out. GO analysis revealed that DEGs were primarily enriched in leukocyte cell–cell adhesion, regulation of leukocyte cell–cell adhesion, and cytokine-mediated signaling pathways (**Fig. 1C**). Similarly, KEGG analysis indicated significant enrichment in pathways including ribosome, NF-kB signaling, and NOD-like receptor signaling (**Fig. 1D**).

**Fig. 1.**
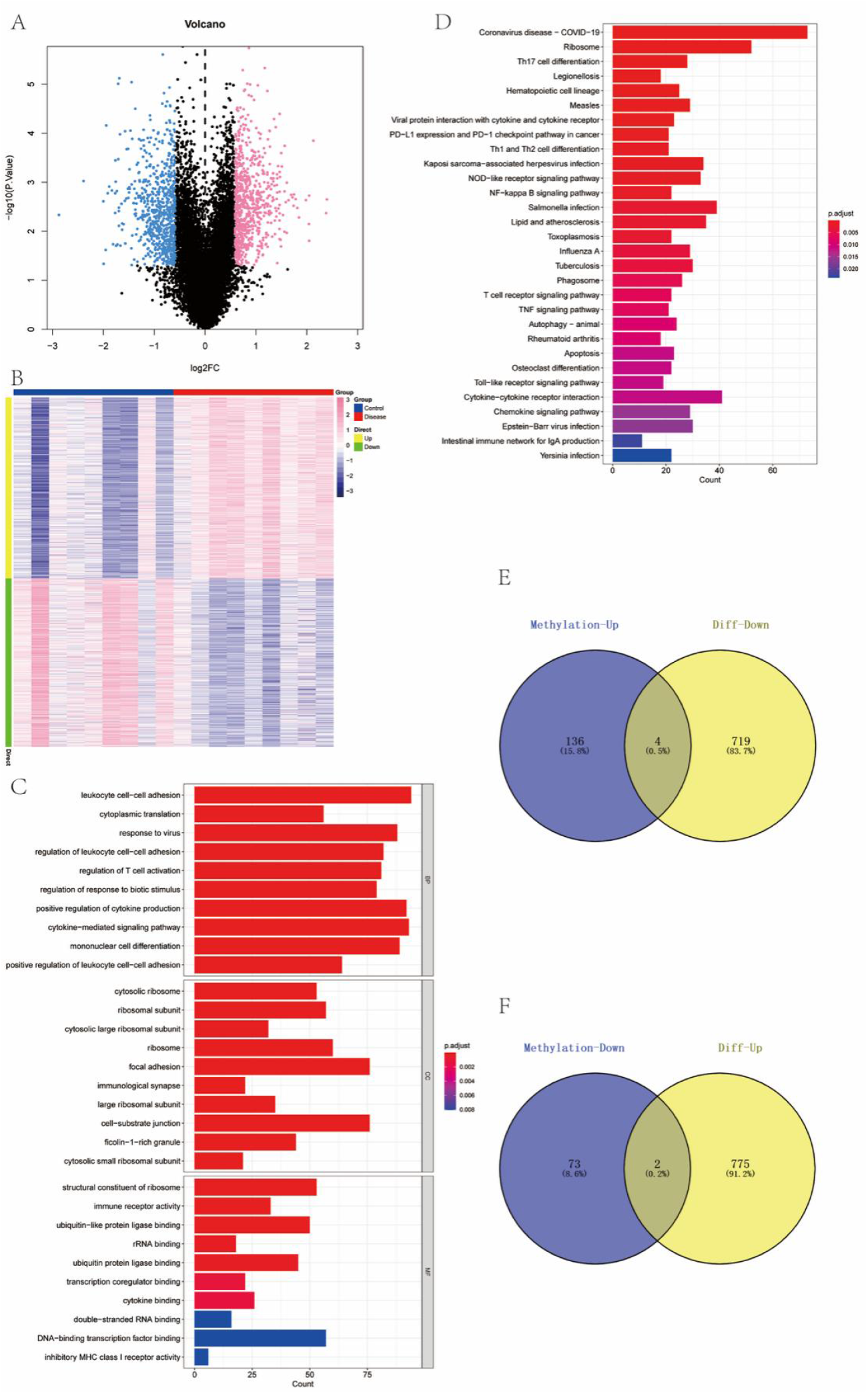
Differential expression and enrichment analyses in APS. A) DEGs in GSE102215 identified by Limma (P < 0.05, |logFC| > 0.585), showing 777 upregulated and 723 downregulated genes; B) complementary visualization of DEGs; C) GO enrichment highlighting leukocyte cell–cell adhesion, regulation of leukocyte cell–cell adhesion, and cytokine-mediated signaling; D) KEGG enrichment highlighting ribosome, NF-kB signaling, and NOD-like receptor signaling; E) Venn intersection integrating DEGs with GSE124565 methylation results; F) six overlapping genes identified.

In parallel, we analyzed methylation data from the GSE124565 dataset, which contained 22 samples, including 12 healthy controls and 10 APS patients, based on the 450K methylation platform. The differential methylation site analysis was performed using ChAMP package, and the screening condition was adj.P.Val < 0.05. There were 91 down-regulated probes (annotated to 75 genes) and 178 up-regulated probes (annotated to 141 genes) (**Supplementary Data**). To integrate expression and methylation results, Venn diagrams were constructed to identify overlap between hypomethylated and differentially expressed genes, yielding 6 intersecting genes (**Fig. 1E–F**).

### 2. Coexpression Network Analysis and Key Gene Identification

To determine the coexpression network of genes in the antiphospholipid syndrome cohort, the WGCNA network was further constructed based on the expression profile data of GSE102215. The soft threshold β was set to 3(**Fig2.A**), and then gene modules were detected based on tom matrix. A total of 5 gene modules were detected in this analysis (**Fig2.B-C**). They were turquoise (3393), blue (1165), black (68), green (115) and red (259), among which turquoise module had the highest correlation with disease (cor=-0.72, p=7e-04). We intersected 3393 module genes in the turquoise module with 6 methylation intersection genes to obtain a total of 2 intersection genes, which will be the key genes in our subsequent studies, namely CCL5 and ITK (**Fig2.D**).

**Fig. 2.**
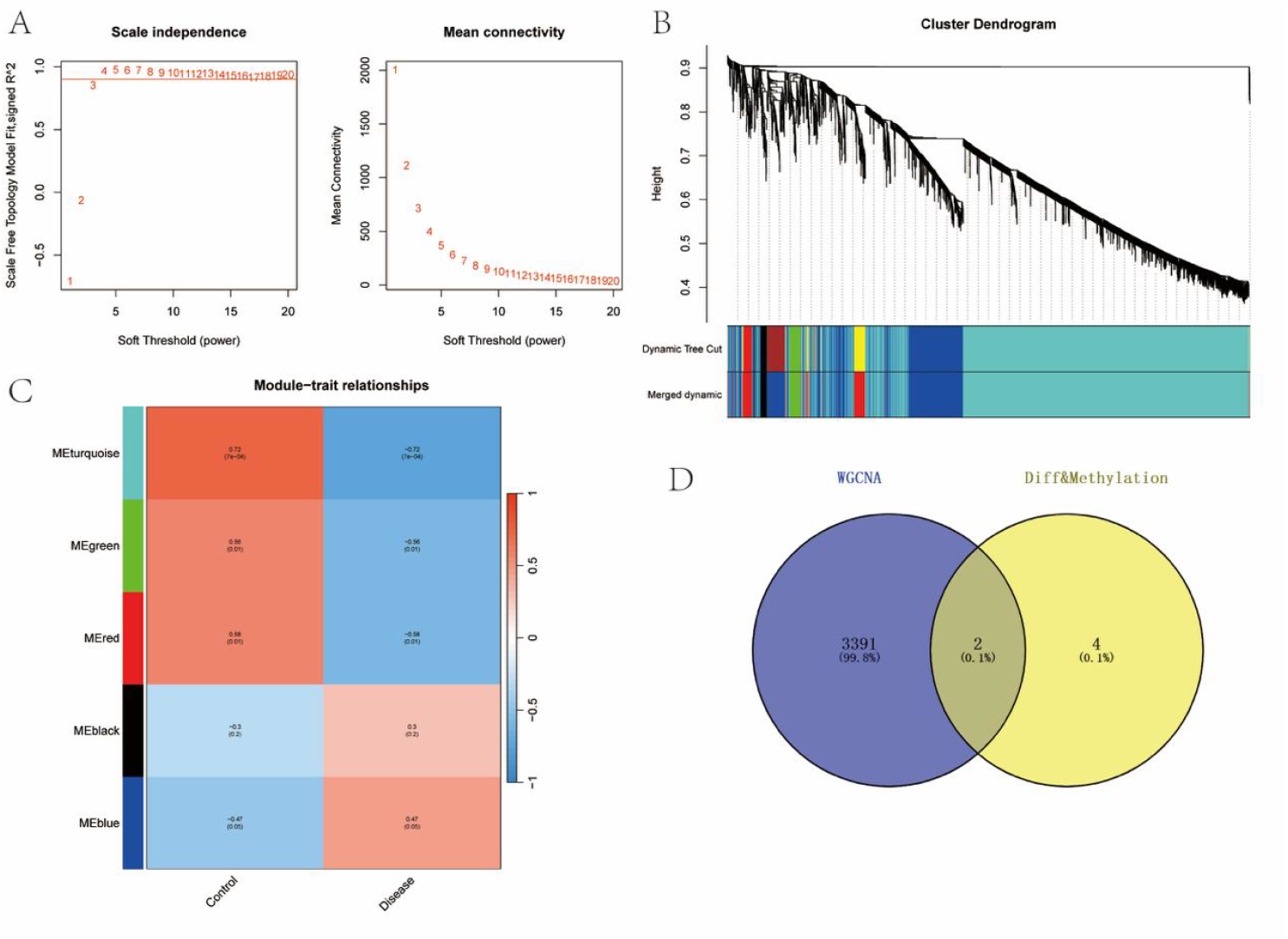
WGCNA module detection and key gene selection. A) Soft-threshold selection with β = 3; B) gene dendrogram and module assignment based on TOM; C) five modules detected, with the turquoise module showing the strongest disease correlation (cor = −0.72, p = 7e−04); D) intersection of turquoise module genes (n = 3,393) with six methylation-based intersections, yielding two key genes, CCL5 and ITK.

### 3. Association of Key Genes with Immune microenvironment

The immune microenvironment is mainly composed of immune-related fibroblasts, immune cells, extracellular matrix, a variety of growth factors, inflammatory factors and special physicochemical characteristics. The immune microenvironment significantly affects the diagnosis, survival outcome and clinical treatment sensitivity of diseases.We obtained the correlation between key genes and different immune factors from the TISIDB database, including immunomodulators, chemokines, and cell receptors (**Fig. 3**). These analyses confirmed that these key genes are potentially related to the immune microenvironment.

**Fig. 3.**
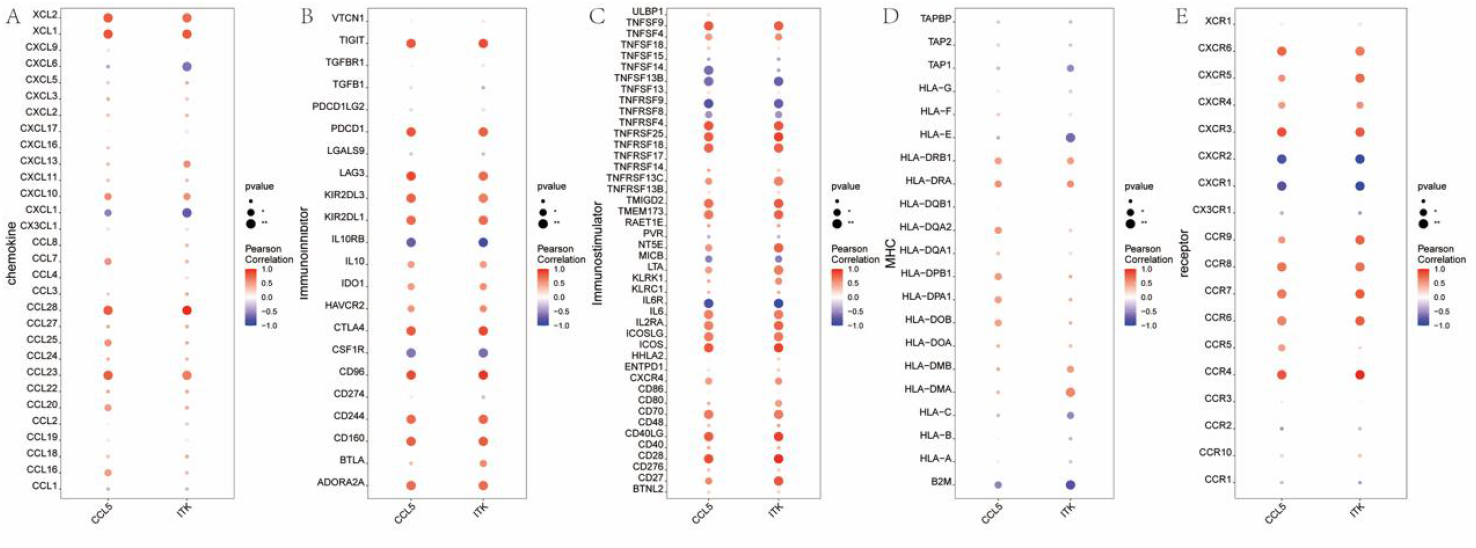
Associations of key genes with immune factors. A) correlations from TISIDB between CCL5 and immune modulators, chemokines, and receptors; B) corresponding correlations for ITK.

### 4. Pathway Enrichment and Functional Analysis

Next, we will study the specific signaling pathways of key gene enrichment and explore the potential molecular mechanisms by which key genes affect the progression of antiphospholipid syndrome. GSEA analysis results showed that CCL5 was mainly enriched in signaling pathways such as Hematopoietic cell lineage, Ribosome, Ribosome biogenesis in eukaryotes (**Fig**.4A-B). In contrast, ITK is mainly enriched in signaling pathways such as Ribosome, Ribosome biogenesis in eukaryotes, and T cell receptor signaling pathway (Fig.4.C-D). GSVA analysis showed that high expression of CCL5 was mainly concentrated in IL6 JAK STAT3 SIGNALING, E2F TARGETS and other signaling pathways (**Fig. 4E**). Similarly, ITK overexpression was enriched in MYC TARGETS V2, E2F TARGETS, and IL2 STAT5 SIGNALING (**Fig. 4F**).

**Fig. 4.**
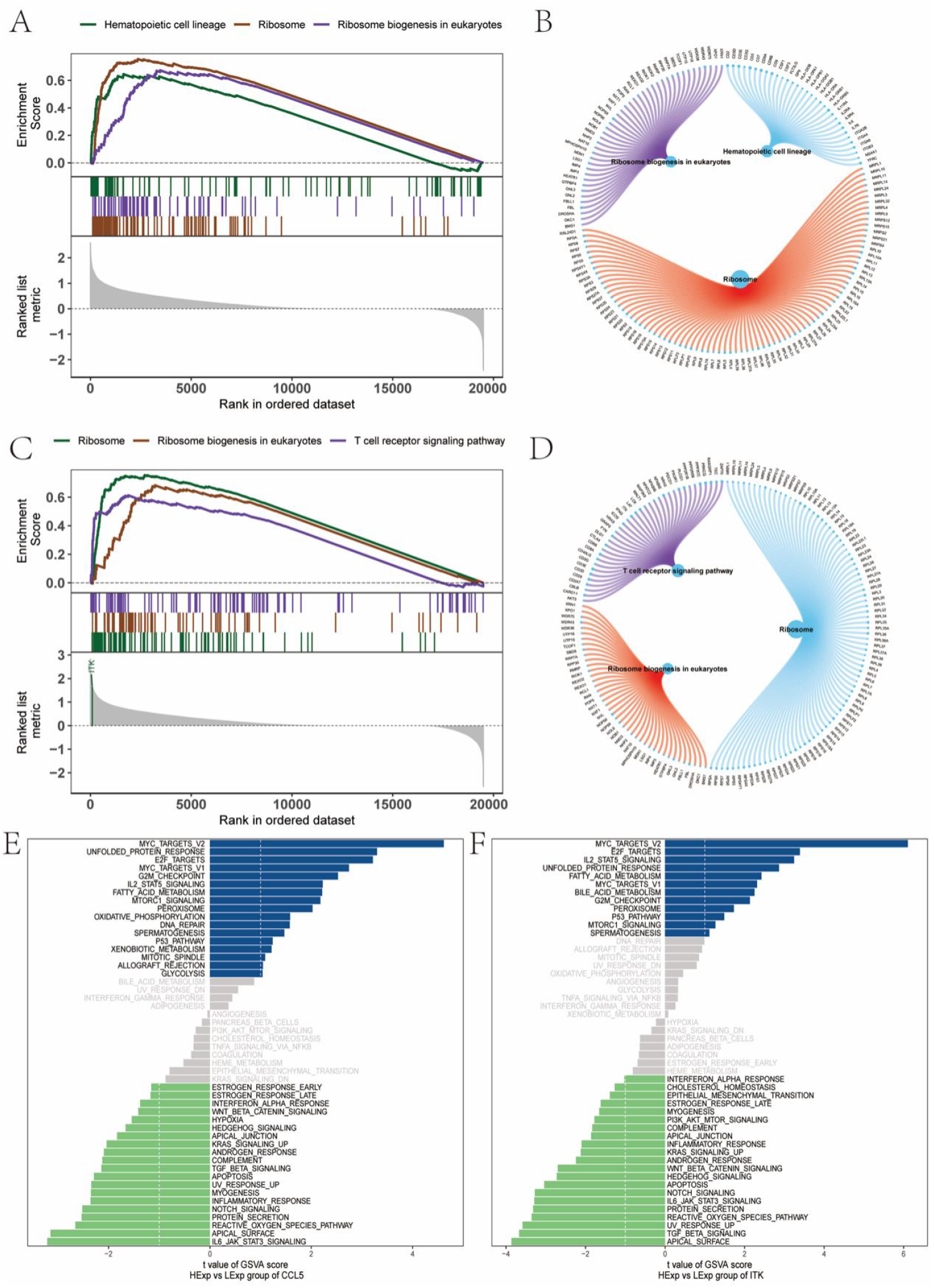
Pathway programs linked to CCL5 and ITK. A) GSEA for CCL5 highlighting hematopoietic cell lineage and ribosome; B) GSEA for CCL5 highlighting ribosome biogenesis in eukaryotes; C) GSEA for ITK highlighting ribosome and ribosome biogenesis in eukaryotes; D) GSEA for ITK highlighting T-cell receptor signaling; E) GSVA showing high CCL5 associated with IL2–STAT5 signaling, MTORC1 signaling, and P53 pathway; F) GSVA showing high ITK associated with IL2–STAT5 signaling, P53 pathway, and MTORC1 signaling.

### 5. Regulatory Network Analysis

To further investigate, we applied these two key genes to the gene set in this analysis and found that they are regulated by common mechanisms such as multiple transcription factors. Therefore, the accumulation recovery curves were used for enrichment analysis of these transcription factors, and the results of Motif-TF annotation and selection analysis of important genes showed that the Motif with the highest standardized enrichment score (NES: 6.56) was cisbp M5814. We demonstrated all the enriched Motifs and corresponding transcription factors of key genes (**Fig. 5A-B**). In addition, reverse prediction using the TargetScanHuman database identified 1,008 miRNAs targeting the two genes, corresponding to 1,104 mRNA–miRNA regulatory pairs, which were visualized in Cytoscape (**Fig. 5C**).

**Fig. 5.**
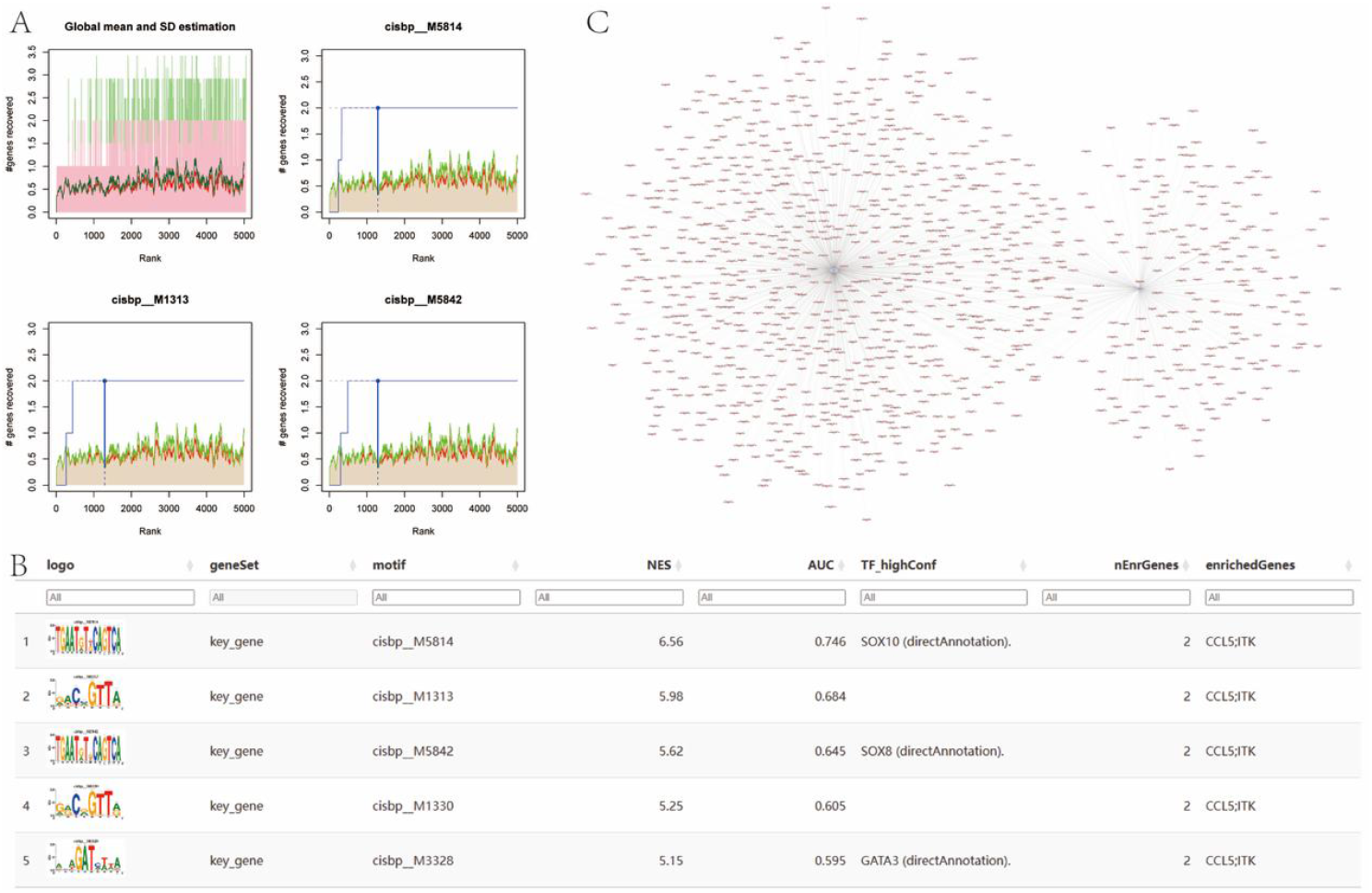
Regulatory networks for CCL5 and ITK. A) transcription factor Motif enrichment with top Motif cisbp__M5814 (NES = 6.56); B) additional enriched Motifs and mapped TFs; C) TargetScanHuman-derived miRNA–mRNA network visualized in Cytoscape (1,008 miRNAs, 1,104 pairs).

### 6. Integration with Disease-Associated Genes and Drug Prediction

Finally, to place the identified key genes in the broader context of APS pathogenesis, we investigated their relationship with known disease-associated genes and potential therapeutic agents. Using the GeneCards database (https://www.genecards.org/), we retrieved APS-related genes and analyzed inter-group expression differences. Analysis of inter-group expression differences of disease genes showed that the expressions of MECP2, CD40LG, AIRE, CDH1 and other genes were different between the two groups of patients (**Fig. 6A**). Furthermore, we analyzed the expression levels of 2 key genes and the top 20 genes of their Relevance score and found that the expression levels of these 2 key genes were significantly correlated with the expression levels of several disease-related genes, and CCL5 and PTEN were significantly negatively correlated (r=-0.624). ITK and CD40LG were significantly positively correlated (r=0.933) (**Fig. 6B**). In addition, differential genes were used for drug prediction through Connectivity Map database, and the results showed that the expression profiles of drug perturbations and disease perturbations were most negatively correlated. It is suggested that drugs can reduce or even reverse the disease state (**Fig. 6.C-F)**.

**Fig. 6.**
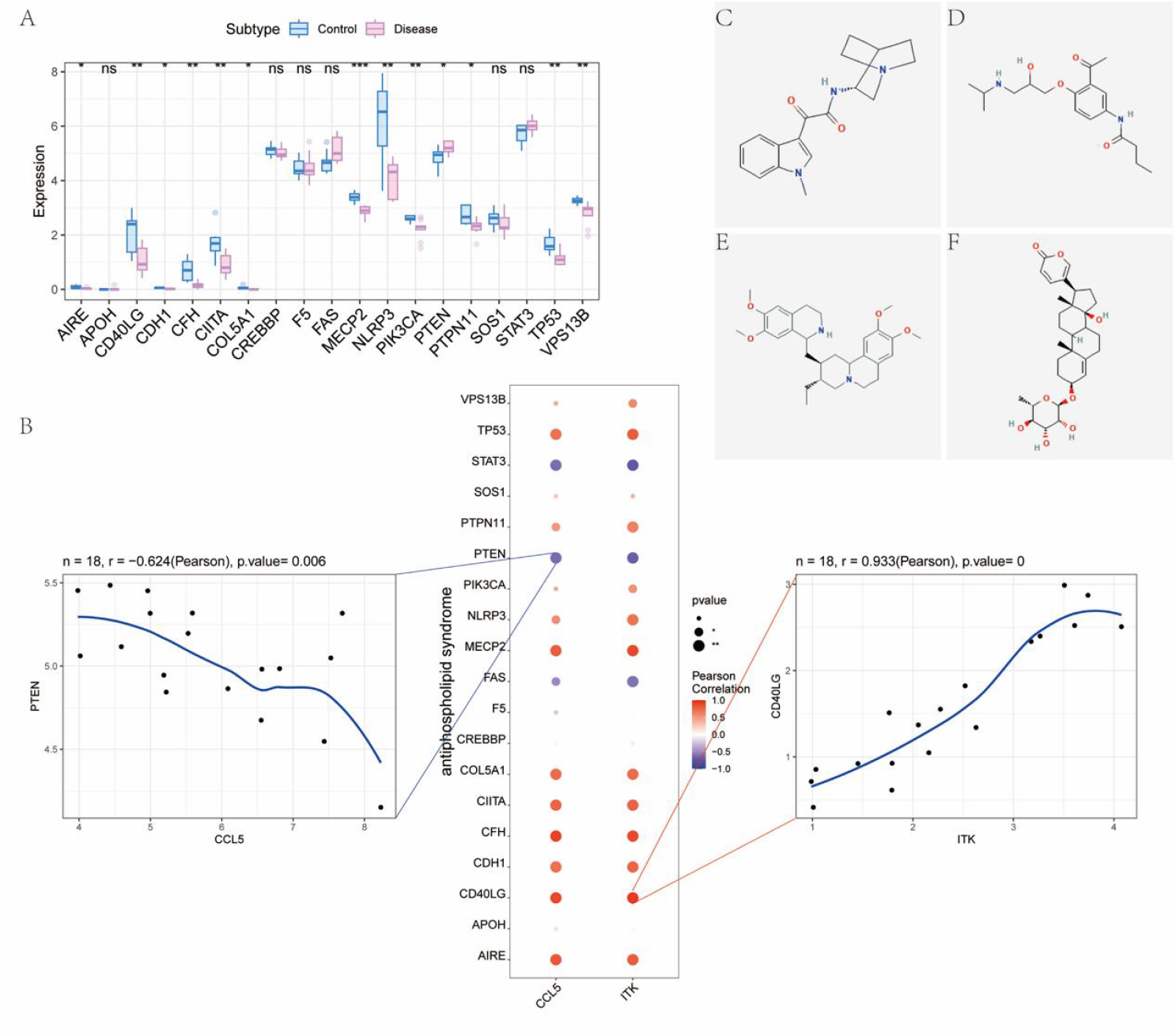
Disease-gene integration and drug predictions. A) differential expression of APS-related genes from GeneCards, including MECP2, CD40LG, AIRE, and CDH1; B) correlations of key genes with disease genes, showing CCL5–PTEN negative correlation (r = −0.624) and ITK–CD40LG positive correlation (r = 0.933); C) Connectivity Map compound with inverse signature, RS-56812; D) acebutolol; E) emetine; F) proscillaridin.

## Discussion

Neutrophils play a key role in the pathogenesis of APS. In this study, we first identified key DEGs of neutrophils in APS using transcriptome, methylation, and WGCNA analyses. Subsequently, we analyzed the relationships between these key DEGs and immune pathways, and we inferred their associations with immune infiltration. We then conducted gene difference analysis for the disease and performed drug prediction using the GeneCards and Connectivity Map databases.

Accordingly, we analyzed neutrophil data in APS from the GSE102215 dataset and identified 1,500 DEGs, including 777 upregulated and 723 downregulated genes. GO and KEGG enrichment analyses indicated strong associations with leukocyte cell–cell adhesion, regulation of leukocyte cell–cell adhesion, cytokine-mediated signaling pathway, ribosome, NF-kB signaling pathway, NOD-like receptor signaling pathway, and other immune response pathways. We then employed a multi-omics WGCNA methodology. A total of five gene modules were detected, among which the turquoise module (3,393 genes) showed the highest correlation with disease. By intersecting the 3,393 turquoise module genes with the six DMGs from the GSE124565 dataset, we identified two key genes associated with neutrophils in APS, namely CCL5 and ITK.

Building on these observations, we note that the roles of CCL5 and ITK in APS have not been investigated previously. CCL5 belongs to the CC chemokine family, also known as RANTES, and is mainly expressed on T lymphocytes and monocytes (Chen et al., 2020; Zaki, 2019). CCL5 protein is secreted primarily by T lymphocytes, macrophages, specific tumor cells, and synovial fibroblasts, and it can activate protein kinase C by binding to receptors CCR1, CCR3, CCR4, and CCR5, with CCR5 showing the strongest affinity, thereby promoting recruitment and migration of diverse immune cells to inflammatory sites (Korbecki et al., 2020). The ITK gene encodes a 72-kDa protein tyrosine kinase that functions in a signal transduction pathway specific to T lymphocytes (Liu et al., 2019). ITK has been reported to differentially regulate T-cell activation, proliferation, and differentiation (Aldinucci et al., 2020), as well as NK cell-mediated cytotoxicity (Sordo-Bahamonde et al., 2020). Fang Zheng et al. found that ITK mutations reduced survival in children with hemophagocytic lymphohistiocytosis after Epstein–Barr virus infection (Chen et al., 2023), suggesting a link to cellular immune dysfunction. Consistent with these observations, our study indicates that CCL5 and ITK are closely related to activation, proliferation, and differentiation of CD8+ T cells and to multiple immunomodulatory factors, which supports their key roles in the immune microenvironment. The observed correlations between key genes and immunomodulators in our study are exploratory in nature and were derived from a pan-cancer context.

At the pathway level, we further observed that CCL5 and ITK were mainly enriched in ribosome, ribosome biogenesis in eukaryotes, IL-2–STAT5 signaling, mTORC1 signaling, and the P53 pathway. These results suggest that the two key genes may influence APS development and progression through these signaling pathways. Among these, existing studies have implicated the mTOR pathway in APS-related thrombosis (Tang et al., 2023). Xie W et al. reported that sirolimus, an mTOR inhibitor, significantly improved thrombocytopenia in patients with APS (Xie et al., 2022). Moreover, the IL-2–STAT5 signaling pathway regulates expression of the transcription factor Foxp3 (Wang & Huang, 2020), and is closely related to differentiation and growth of CD4+ and CD25+ Treg cells and to immune regulation (Luckheeram et al., 2012). Dal Ben ER et al. found that circulating CD4+ and CD25+ Treg cells are reduced in APS compared with healthy individuals, thereby promoting APS development (Yan et al., 2022). By contrast, ribosome, ribosome biogenesis in eukaryotes, and the P53 pathway have not been studied in combination with APS. In addition, our findings suggest that CCL5 and ITK can be regulated by common mechanisms involving multiple transcription factors. In our study, we found three key transcription factors: SOX10, SOX8 and GATA3.The SOX gene encodes transcription factors involved in embryogenesis and cell differentiation. Specifically, the transcription factor SOX10 plays a critical role in the differentiation of neural crest precursors into the melanocytic lineage (Qi et al., 2022). Sheera R Rosenbaum et al. demonstrated that genetic ablation of SOX10 enhanced the susceptibility of melanoma cells to CD8+ T cell-mediated killing and cell death induction by TNFα or IFNγ. In SOX10-deficient cells, cytokine-induced cell death exhibited characteristics of caspase-dependent pyroptosis an inflammatory form of cell death with the potential to amplify immune responses (Rosenbaum et al., 2024). SOX8 belongs to the Sox family and is closely related to SOX9 and SOX10. During prenatal development, SOX8 is expressed in multiple tissues derived from ectoderm, endoderm, and mesoderm, and participates in organogenesis and differentiation processes. Its expression is observed in various important cell types, such as glial cells, satellite cells, and chondrocytes. SOX8 plays a more prominent role in late development and adult tissues. Genetic variations in SOX8 have been associated with multiple sclerosis and familial essential tremor, while alterations in SOX8 are also linked to poor cancer prognosis and infertility (González Alvarado & Aprato, 2025). GATA binding protein 3 (GATA3) is a zinc-finger pioneer transcription factor involved in diverse biological processes. It regulates gene expression by binding nucleosomal DNA and facilitating chromatin remodeling, with its activity being modulated by post-translational modifications. During development, GATA3 plays a key role in cell differentiation. Mutations in GATA3 are associated with breast and bladder cancers, and have significant implications for immune status and therapeutic response (Zhang et al., 2022).These transcription factors directly or indirectly influence the tumor immune microenvironment and modulate immune response efficiency by regulating cell differentiation and gene expression programs.

Complementing these pathway insights, our study also showed that CCL5 and ITK were significantly associated with expression of multiple genes related to APS. There was a significant negative correlation between CCL5 and PTEN, and a significant positive correlation between ITK and CD40LG. PTEN encodes a phosphatase for PIP3, the lipid product of PI3-kinase, and directly counteracts activation of the oncogenic PI3K/AKT/mTOR pathway (He et al., 2021). Rubén Armañanzas et al. identified a set of 150 genes that distinguishes SLE and PAPS patients from healthy individuals (Armananzas et al., 2009). Functional characterization indicated interferon regulation and also revealed other altered regulatory pathways, including PTEN, TNF, and BCL-2, in SLE and PAPS. These observations support a potential link between PTEN alteration and APS onset. The X-linked gene CD40LG encodes CD40L, a T-cell coactivator receptor whose expression is controlled by NFAT transcription. CD40L-deficient patients exhibit impaired immunoglobulin class switching and cellular immunodeficiency, whereas mice overexpressing CD40L develop high autoantibody titers and chronic inflammation (Meryem et al., 2025). Anh Nguyen Lien Phan et al. reported that patients with X-linked hyper-IgM syndrome caused by CD40LG mutations developed APS (Phan et al., 2021; Vastrad et al., 2025). These results suggest that CD40LG gene mutation may be associated with APS onset.

Furthermore, we performed drug prediction analysis by submitting the differentially expressed genes to the Connectivity Map (CMap) database. The results revealed that the gene expression changes induced by perturbations of drugs such as RS-56812, Acebutolol, Emetine, and Proscillaridin showed a significant negative correlation with the expression pattern observed in the disease state, suggesting that these compounds may have the potential to reverse disease-associated molecular phenotypes.RS-56812, a 5-HT3 receptor antagonist, has been reported to exert potential therapeutic effects on cognitive decline by Terry AV Jr et al (Cowen & Sherwood, 2013). Acebutolol is a cardioselective beta-adrenoceptor blocker with partial agonist and membrane-stabilizing activity for patients with hypertension and arrhythmias (Čižmáriková et al., 2019). Emetine is an effective treatment for amebic liver abscess, and it has recently demonstrated antiviral efficacy against RNA and DNA viruses (Islam et al., 2024; Panwar et al., 2020). Proscillaridin is a cardiac glycoside used as a Na^+^/K^+^ pump inhibitor in heart failure. Recently, Da Costa EM et al. found that proscillaridin induced MYC degradation, epigenetic reprogramming, and differentiation of leukemia cells through loss of lysine acetylation, specifically targeting MYC-overexpressing leukemia cells and leukemia stem cells (Poohadsuan et al., 2023). These agents have not been studied in combination with APS in previous literature, and our results suggest that they may have potential therapeutic value for APS.It should be noted, however, that the drug predictions generated in this study represent preliminary hypotheses based on public database queries. Their actual biological effects are likely highly context-dependent, influenced by factors such as cell type, dosage, and model systems. Moreover, the reproducibility of these predictions across independent datasets remains to be further validated through subsequent experimental studies.

Finally, it must be noted that the present study has limitations. Correlations between key genes and disease-associated genes were calculated using Pearson’s correlation. P-values were adjusted for multiple testing using the Benjamini–Hochberg false discovery rate (FDR) method. Only correlations with an FDR < 0.05 were considered statistically significant. We emphasize that these correlations do not imply causation.In subsequent experiments, we will collect fresh blood samples to assess differences in gene expression and methylation levels of neutrophils in APS, thereby addressing limitations of the current analysis. In addition, we will further validate the APS animal model and conduct experimental evaluation of the predicted drugs.

## Conclusion

This work integrates transcriptome and epigenome profiles to define a neutrophil-centered molecular signature in APS. By combining Limma-based differential expression with WGCNA and methylation integration, we prioritized two key genes, CCL5 and ITK, that aligned with immune infiltration patterns, including higher follicular helper T cells, activated dendritic cells, and neutrophils in APS cases. Consistent pathway signals from GSEA and GSVA, including IL-2–STAT5, mTORC1, P53, and ribosome-related programs, point to coordinated transcriptional programs that may shape neutrophil–lymphocyte cross-talk and thrombo-inflammatory activity in APS. Moreover, correlations of CCL5 with PTEN and of ITK with CD40LG, together with Motif and miRNA regulatory analyses, support a model in which these genes participate in broader immune regulatory circuits relevant to APS pathobiology. Finally, Connectivity Map outputs nominate small-molecule candidates, such as RS-56812, acebutolol, emetine, and proscillaridin, that exhibit inverse transcriptomic signatures and therefore merit experimental assessment as potential modulators of APS-related pathways.

## Declarations

### Data availability

All data used in this study were retrieved from public resources. Gene expression and methylation datasets were obtained from GEO (GSE102215 and GSE124565). Auxiliary resources, including GeneCards, Connectivity Map, TISIDB, and MSigDB, are publicly accessible. Relevant processed data or analysis files that are not already deposited publicly are available from the corresponding author on reasonable request. Requests for materials, analysis scripts, or additional clarifications should be directed to the corresponding author.

### Author contributions

MC conceived and designed the study, acquired and curated the data, performed the analyses, developed the software, generated the figures, and wrote the first draft of the manuscript. YLi supervised the work, provided resources, critically revised the manuscript for important intellectual content, and approved the final version. Both authors contributed to the interpretation of results, approved the final manuscript, and agree to be accountable for all aspects of the work.

